# DMT1 knockout abolishes ferroptosis induced mitochondrial dysfunction in *C. elegans* amyloid β proteotoxicity

**DOI:** 10.1101/2024.08.08.607074

**Authors:** Wilson Peng, Kaitlin B Chung, B Paige Lawrence, M Kerry O’Banion, Robert T Dirksen, Andrew P Wojtovich, John O Onukwufor

## Abstract

Iron is critical for neuronal activity and metabolism, and iron dysregulation alters these functions in age-related neurodegenerative disorders, such as Alzheimer’s disease (AD). AD is a chronic neurodegenerative disease characterized by progressive neuronal dysfunction, memory loss and decreased cognitive function. AD patients exhibit elevated iron levels in the brain compared to age-matched non-AD individuals. However, the degree to which iron overload contributes to AD pathogenesis is unclear. Here, we evaluated the involvement of ferroptosis, an iron-dependent cell death process, in mediating AD-like pathologies in *C. elegans*. Results showed that iron accumulation occurred prior to the loss of neuronal function as worms age. In addition, energetic imbalance was an early event in iron-induced loss of neuronal function. Furthermore, the loss of neuronal function was, in part, due to increased mitochondrial reactive oxygen species mediated oxidative damage, ultimately resulting in ferroptotic cell death. The mitochondrial redox environment and ferroptosis were modulated by pharmacologic processes that exacerbate or abolish iron accumulation both in wild-type worms and worms with increased levels of neuronal amyloid beta (Aβ). However, neuronal Aβ worms were more sensitive to ferroptosis-mediated neuronal loss, and this increased toxicity was ameliorated by limiting the uptake of ferrous iron through knockout of divalent metal transporter 1 (DMT1). In addition, DMT1 knockout completely suppressed phenotypic measures of Aβ toxicity with age. Overall, our findings suggest that iron-induced ferroptosis alters the mitochondrial redox environment to drive oxidative damage when neuronal Aβ is overexpressed. DMT1 knockout abolishes neuronal Aβ−associated pathologies by reducing neuronal iron uptake.

**Highlights:**

1. Energetic imbalance is an early event in iron-induced loss of neuronal function
2. Neuronal Aβ increases susceptibility to ferroptosis mediated oxidative damage
3. Divalent metal transporter 1 knockout protects against iron-induced oxidative damage and ferroptosis

**Graphical Abstract:** 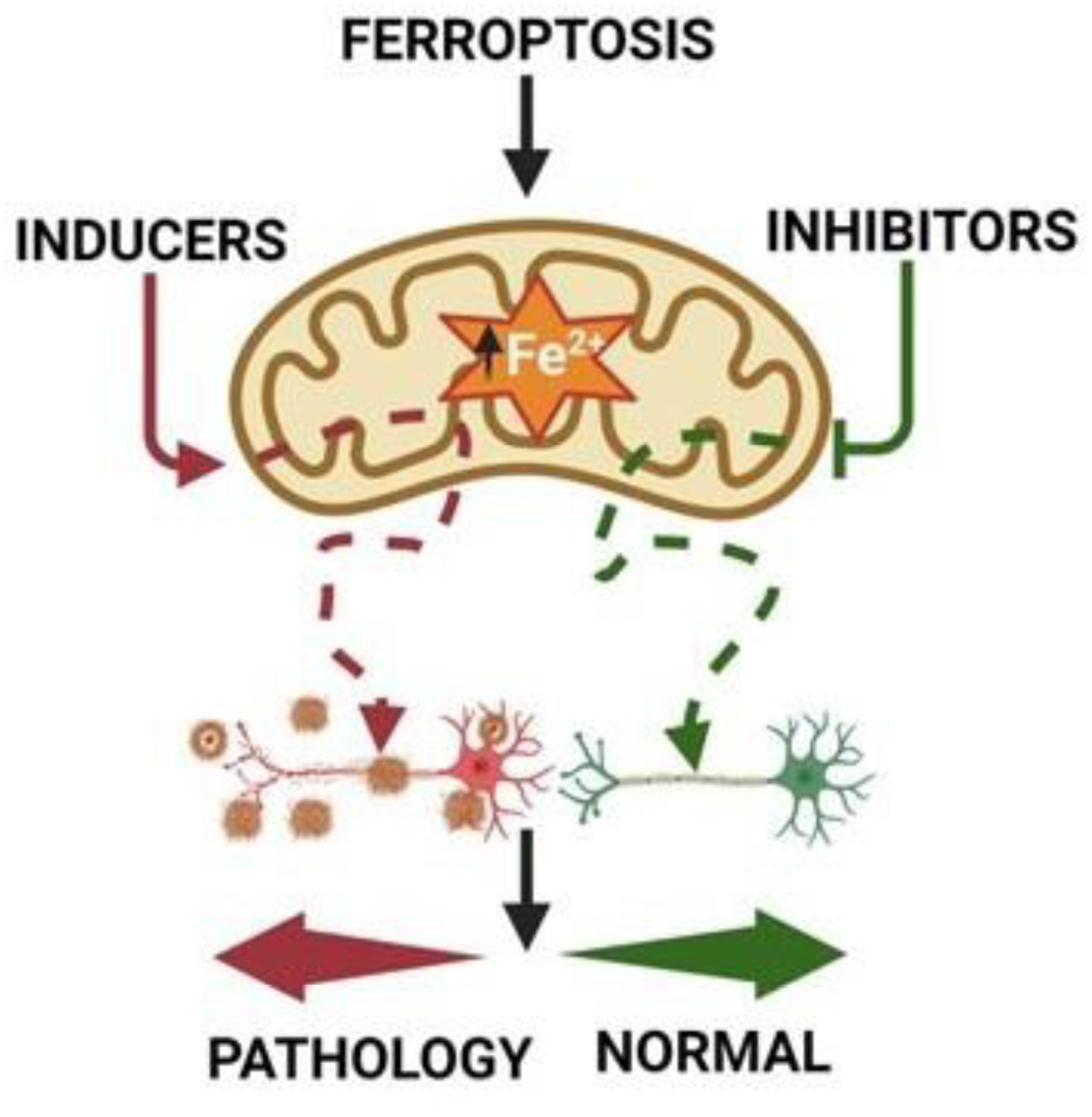

## 1. Introduction

Iron is essential for cellular function, where its tight regulation aids in neurotransmitter biosynthesis, myelination, and energy homeostasis[1–3]. Cellular iron levels are controlled by iron regulatory proteins. These include divalent metal transporter 1 (DMT1), which facilitates cytosolic ferrous iron uptake, and ferritin, which enables the storage of ferrous iron in the form of ferric iron, and ferroportin, which exports ferrous iron out of the cytosolic environment[3, 4]. Free ferrous iron not stored or exported from the cell is utilized for a variety of cellular functions. Mitochondria use ferrous iron for heme biosynthesis, iron-sulfur cluster formation and mitochondrial electron transport chain (ETC) activity for energy production[3, 5, 6]. However, as organisms age, the efficiency of iron regulatory proteins to sequester and control cytosolic free ferrous iron declines, leading to excess free ferrous iron levels that impair mitochondrial function[7, 8].

The impairment of mitochondrial activity by excess ferrous iron results in lower cellular ATP production and higher mitochondrial reactive oxygen species (ROS) production[5, 9]. Mitochondrial ROS, when produced in small quantities, are important for cellular signaling[10–12]. However, large increases in mitochondrial ROS, as occurs with iron toxicity, trigger oxidative stress and damage to membrane proteins and lipids[9, 11]. Mitochondrial ROS can also react with ferrous iron to produce potent reactive free radicals that cause cellular death by ferroptosis[3]. Ferroptosis is an iron-dependent programmed cell death mechanism mediated by ROS induced lipid peroxidation[13, 14]. Ferroptosis is associated with many age-related disorders, such as Alzheimer’s disease (AD)[3, 6]. However, the underlaying cellular and molecular mechanisms, as well as the degree to which iron-induced ferroptosis contributes to AD pathogenesis remains unknown.

Model organisms offer a tractable system to interrogate causal relationships in order to fine-tune our mechanistic understanding of how iron dysregulation contributes to AD pathogenesis. *C. elegans* is an excellent model to investigate the cellular and molecular mechanisms of mitochondrial iron dysregulation in AD[15, 16]. In *C. elegans,* movement disorders correlate with a decline in neural function, while swimming rates provide a reliable read out of energetic output. Both of these phenotypes correlates with the observed pathology in AD patients[17, 18].

Therefore, we used wild-type *C. elegans,* as well as *C. elegans* overexpressing neuronal Aβ, to elucidate how iron dysregulation contributes to altered physiology. We found that energetic imbalance measured by impaired swimming rate and mitochondrial dysfunction represent early events in iron-induced toxicity. Iron toxicity was facilitated by ROS induced oxidative damage that promotes ferroptosis. Pharmacologic and genetic methods of inhibiting or promoting ferroptosis either exacerbated or abolished, respectively, ferroptotic pathologies. Furthermore, worms with over expression of neuronal Aβ were more sensitive to iron induced toxicity than age-matched wild-type worms and knockout of DMT1 mitigated this enhanced iron-induced pathology of neuronal Aβ. These findings suggest that enhanced iron toxicity promotes Aβ pathology and that iron regulatory proteins such as DMT1 represent potential therapeutic targets to mitigate Aβ-mediated toxicity.

## 2. Materials and Methods

### Worm maintenance and strains

*C. elegans* were maintained on OP50 bacterial lawns on nematode growth media (NGM). The following strains were used in this investigation: wildtype [N2]; CL2355 [*smg-1(cc546) dvls50 I*]; RB1074 [*smf-3(ok1035) IV*], and OOJ1 [*smg-1(cc546) dvls50 I*; *smf-3(ok1035) IV*]. CL2355 and OOJ1 were maintained at 16°C, and all other strains were maintained at 20°C. All the strains used in this study, with the exception of the OOJ1 strain generated in house, were provided through the *Caenorhabditis* Genetics Center (CGC).

### Mitochondrial isolation

Mitochondria were isolated by differential centrifugation as previously described[12]. In brief, approximately 0.25 million synchronized L4 worms were grown on HB101 with iron (0 or 35 µM) for 3 days. Worms were then rinsed with M9 media (22 mM KH_2_PO_4_, 42 mM Na_2_HPO_4_, 86 mM NaCl, 1 mM MgSO_4_, pH 7) before transferring to mitochondrial isolation media (220 mM mannitol, 70 mM sucrose, 5 mM MOPS, 2 mM EGTA, pH 7.3 at 4°C). Using pure sea sand in an ice-cold mortar, worms were crushed followed by a Dounce homogenization. The homogenate was then passed through a series of differential centrifugations using mitochondrial respiratory media (220 mM mannitol, 70 mM sucrose, 5 mM MOPS, 2 mM EGTA, 0.04% BSA, pH 7.3 at 4°C) to enrich the mitochondrial preparation. The protein concentration of the mitochondrial preparation was determined using the Folin-phenol method. Mitochondrial respiration and superoxide measurements were conducted using freshly isolated mitochondria. Mitochondrial enzyme activity was conducted using frozen mitochondria within 2 weeks of isolation.

### Mitochondrial respiration

Freshly isolated mitochondria were used to measure mitochondrial respiration using a Clark-type O_2_ electrode (Hansatech Instruments, UK) as described previously[12]. In brief, after calibration of the electrode, mitochondria (1 mg/ml) suspended in mitochondrial respiration buffer (120 mM KCl, 25 mM sucrose, 5 mM MgCl_2_, 5 mM KH_2_PO_4_, 1 mM EGTA, 10 mM HEPES pH 7.3) were loaded in the chamber. Mitochondrial complex I powered respiration was measured using complex I-linked substrates, 2.5 mM malate plus 5 mM glutamate. The addition of ADP (0.4 mM) was used to drive state 3 respiration followed by the addition of oligomycin (1 µg/ml) to generate state 4 respiration. All substrates were added to the chamber via a syringe port.

### Mitochondrial enzyme activity

Complex I and citrate synthase activities were assessed by spectrophotometric methods following permeabilization of isolated mitochondria (1 mg/ml) with three bouts of freeze-thaw[12]. Citrate synthase activity was measured as the rate of DTNB-coenzyme A formation with an extinction coefficient of 13600 M^-1^ at 412 nm. Complex I activity was determined as the rotenone-sensitive rate of NADH oxidation with an extinction coefficient of 6180 M^-1^ at 340 nm.

### Mitochondrial superoxide measurement

Superoxide production from freshly isolated mitochondria was assessed using 2-hydroxyethidium (2-OHE^+^)[19]. In brief, proteins were precipitated using 200 mM HClO_4_/MeOH and removed via centrifugation followed by the addition to the supernatant of an equal volume of phosphate buffer (1 M, pH 2.6). Samples were then filtered and separated using a polar-RP column (Phenomenex, 150 x 2 mm; 4 µm) on an HPLC (Shimadzu) with fluorescence detection (RF-20A). Prior to sample analysis, a standard curve was generated using purified 2-OHE^+^. The HPLC protocol included mobile phase A (A: 10% ACN, 0.1 % TFA) and mobile phase B (60% ACN, 0.1 % TFA) consisting of the following gradient: 0 min, 40% B, 5 min, 40% B; 25 min, 100% B; 30 min, 100% B; 35 min 40% B; 40 min, 40% B. Samples were quantified using Lab Solutions (Shimadzu).

### Worm paralysis assessment

Synchronized L4 worms were individually transferred to a seeded plate containing iron or drug every 24 h until the end of the trial (duration of trials indicated in respective figures). Paralysis, defined as the inability to move upon mechanical stimulation, was scored every 24 h[7].

### Worm swimming rate

Synchronized L4 worms were individually transferred to a seeded plate containing iron or drug every 24 h until the end of the trial (duration of trials indicated in respective figures). Non-paralyzed worms were then individually transferred to an unseeded plate containing 100 µl of M9. An acclimatization period of 30 s was included before the assessment of swimming rate calculated over 15 s.

### Lipid peroxidation measurement

*In vivo* quantification of lipid peroxidation in individual anesthetized live worms was performed using the BODIPY 581/591 C11[20]. The degree of lipid ROS production was assessed from a shift in BODIPY 581/591 C11 fluorescence emission peak from 590 to 510 nm due to the oxidation of the polyunsaturated butadienyl portion of C11-BODIPY[20].

### Confocal imaging

Synchronized L4 worms were grown on a plate containing iron (0 or 35 µM) for 5 days. Worms were then transferred to a seeded plate containing 1.25 µM BODIPY 581/591 C11 for 60 min and then anesthetized on a 2% agarose pad containing 0.1% tetramisole. Worms were imaged using a Nikon AXR confocal microscope equipped with a 40x water objective. All imaged data were analyzed using ImageJ software.

### Elemental analyses

Synchronized L4 worms (0.25 million) were transferred to a seeded plate containing iron (0 or 35 µM) for 3 days. For whole worm elemental analysis, worms were harvested by first washing in M9 media with the worm pellet collected via centrifugation and then resuspended in 1 mL M9 volume. For mitochondrial elemental analysis, mitochondria were isolated (see mitochondrial isolation procedure) and then suspended in mitochondrial buffer. Whole worm and mitochondrial, samples were weighed before digestion. To digest samples, 1mL 69 % HNO_3_ and 0.5mL HCl (36%) was added at 100°C for 60 min. Following digestion, samples were cooled, and double deionized water was added up to 10 ml total volume. Samples were analyzed using the following detection limits in ppb (ng/ml): Fe 0.123, Cu 0.0009, Zn 0.0241, Mn 0.00136, and Ca 3.688. Samples were analyzed with ICP-Mass Spectrometry (Perkin Elmer 2000C) and data were normalized to the initial weight of the samples in grams.

### Statistical analyses

All statistical analyses were performed using GraphPad^TM^ Prism v10 (GraphPad Software, San Diego, CA, USA). Data were first subjected to normality and homogeneity variance testing. All data in this study passed the test. Data were then analyzed using either an unpaired, two-tailed student’s t-test, or one or two-way ANOVA with post hoc multiple comparison correction. Means were considered significantly different when the p-value was < 0.05.

## 3. Results

### 3A. Iron toxicity impairs worm physiologic function

Previous studies found that iron toxicity increases worm paralysis with age[7]. Here, we confirmed these findings and established an effective concentration of iron that produced a 50% increase in paralysis after 10 days of exposure at 20°C (Fig. 1A and B). Using 35 µM iron, the lowest effective concentration observed at day 6 at 20°C (Fig. 1B), we determined the impact of iron exposure on two key physiological readouts of neuronal function at 25°C: 1) paralysis and 2) energetic output assessed from swimming rate. These analyses revealed an age-dependent increase in paralysis with a marked difference between control and iron treated worms starting at day 3 of iron exposure (Fig. 1D). In contrast, a significant reduction in swimming rate was observed as early as one day after iron exposure (Fig. 1E), consistent with changes in swimming rate being a more sensitive index of iron toxicity onset. These results suggest that energetic failure and reduced neuronal function represent likely downstream events of iron toxicity that initially lead to reduced swimming rate, and eventually, paralysis.

**Figure 1:**
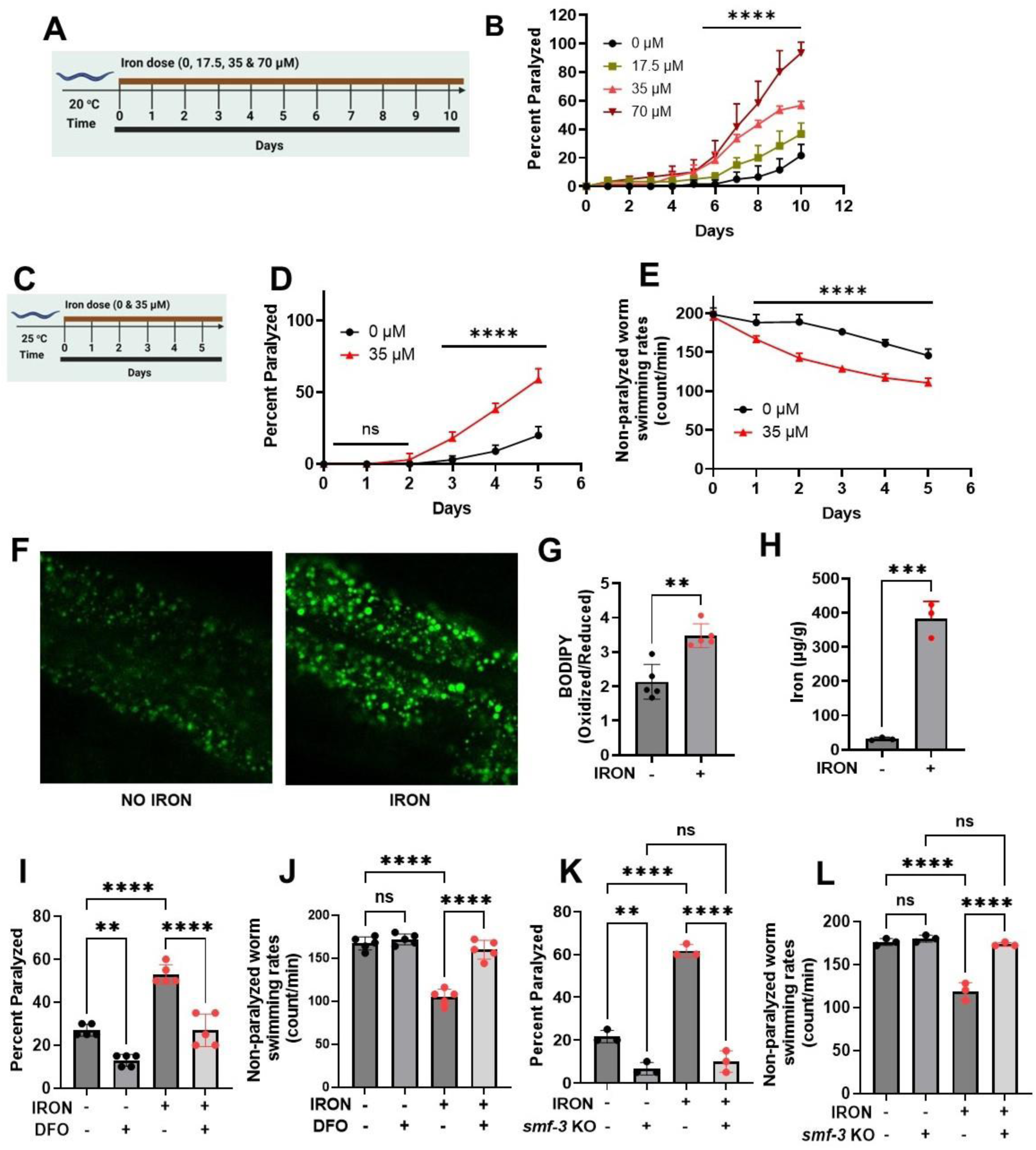
Iron toxicity alters worm physiologic function: **A)**. Experimental layout showing 10 days of worm exposure to different iron doses (0, 17.5, 35, and 70 µM) at 20 °C. **B)**. Dose-and time-dependent effects of iron exposure on worm paralysis at 20 °C. Staged L4 worms were transferred to plates containing iron (0, 17.5, 35, and 70 µM). Worms were then transferred every 24 h for 10 days. Paralysis (e.g., inability to move upon stimulation) was scored every 24 h for 10 days. Data are mean ±SEM, N=3 independent biological replicates (where one biological replicate contains 20 worms per plate). ****p<0.0001, two-way ANOVA, Tukey post hoc test. **C)**. Experimental layout showing 5 days of worm exposure to iron (0 and 35 µM) at 25 °C. **D)**. Time course of iron exposure on worm paralysis at 25 °C. Staged L4 worms were transferred to plates containing iron (0 or 35 µM). Paralysis was scored every 24 h for 5 days. Data are mean ±SEM, N=5 independent biological replicates (where one biological replicate contains 20 worms per plate). ns not significant, ****p<0.0001, one-way ANOVA, Tukey post hoc test. **E)**. Iron reduced non-paralyzed worm swimming rates at 25 °C. Staged L4 worms were transferred to plate containing iron (0 or 35 µM). Worms were then transferred every 24 h for 5 days. Non-paralyzed worms were individually transferred to plate containing 100 µl of buffer. After 30 seconds of equilibration, swimming rates were collected for 15 seconds. Data are mean ±SEM N=5 independent replicates (where 4 independent worm count constitute an N). ****p<0.0001, one-way ANOVA, Tukey post hoc test. Iron toxicity increases whole worm lipid peroxidation. **F)** Confocal Image and **G)** Quantification. Staged L4 worms were transferred to plate containing iron (0 or 35 µM). Worms were then transferred every 24 h for 5 days. Then worms were transferred to plate containing 1.25 µM BODIPY for 60 min and prep for confocal imaging. Image scale 30 µm. Data are mean ±SEM N=5 independent replicates, **p=0.001, one-way ANOVA, Tukey post hoc test. **H)**. Total iron was measured in worms using ICP-MS where dark circle (no iron) and red triangle(iron) bars treated with 35 µM iron. Data are mean ±SEM, N=4 independent biological replicates. ****p<0.0001, Unpaired t test. **I)**. Effects of Deferoxamine (DFO) on iron-induced paralysis. Staged L4 worms were transferred to plates containing 0 µM iron, 0 µM iron + 100 µM DFO 35 µM iron and 35 µM iron + 100 µM DFO. Paralysis was scored every 24 h for 5 days. Data are mean ±SEM, N=5 independent biological replicates (where one biological replicate contains 20 worms per plate). ns not significant, ****p<0.0001, one-way ANOVA, Tukey post hoc test. **J)**. Deferoxamine restored iron-induced impairment of non-paralyzed worm swimming rates. Staged L4 worms were transferred to plates containing 0 µM iron, 0 µM iron + 100 µM DFO 35 µM iron and 35 µM iron + 100 µM DFO . Non-paralyzed worms were individually transferred to plate containing 100 µl of buffer. After 30 seconds of equilibration, swimming rates were collected for 15 seconds on day 5. Data are mean ±SEM N=5 independent replicates. ns not significant, ****p<0.0001, one-way ANOVA, Tukey post hoc test. **K)**. DMT1 (*smf-3*) knock out abolished iron toxicity mediated worm paralysis. Staged L4 worms were transferred to plates containing iron WT, *smf-3* KO, WT + iron (35 µM) and *smf-3* KO + iron (35 µM). Paralysis was scored every 24 h for 5 days. Data are mean ±SEM, N=3 independent biological replicates (where one biological replicate contains 20 worms per plate). *p<0.05, ****p<0.0001, one-way ANOVA, Tukey post hoc test. **L)**. DMT1 (*smf-3*) knock out shielded worms from iron-induced reduction of non-paralyzed worm swimming rates. Staged L4 worms were transferred to plates containing iron WT, *smf-3* KO, WT + iron (35 µM) and *smf-3* KO + iron (35 µM). Non-paralyzed worms were individually transferred to plate containing 100 µl of buffer. After 30 seconds of equilibration, swimming rates were collected for 15 seconds. Data are mean ±SEM N=3 independent replicates (where 4 independent worm count constitute an N). ns not significant, ***p=0.0001, ****p<0.0001, one-way ANOVA, Tukey post hoc test.

Iron toxicity could be mediated by lipid peroxidation. Therefore, we measured whole worm lipid peroxidation in both control and iron-treated worms using BODIPY C11, a probe used to assess lipid oxidization mediated by ferrous iron[20]. Iron-treated worms exhibited higher levels of oxidized lipids relative to non-treated age-matched worms (Fig. 1F&G), potentially due to increased iron overload. To directly test this idea, we used ICP-MS to quantify total iron load in control and iron-treated worms. A significant increase in total iron content was observed in iron-treated worms (Fig. 1H), consistent with iron exposure leading to iron overload and toxicity. To further confirm this connection, we tested the ability of exposure to a pharmacologic iron chelator (deferoxamine or DFO) to reverse the effects of iron[21]. These studies revealed that DFO exposure prevented both iron mediated worm paralysis (Fig. 1I) and reduced swimming rate (Fig. 1J). However, DFO does not discriminate between ferrous and ferric iron. In order to determine the iron species that is responsible for altered physiologic processes, we used genetic worms deficient in divalent metal transporter 1 (DMT1 [*smf-3*]), which specifically facilitates ferrous iron uptake[3, 22–24]. We hypothesized that *smf-3*-deficient worms will exhibit reduced ferrous iron uptake, and thus, reduced iron-dependent toxicity. Consistent with these expectations, *smf-3* mutant worms lacked iron dependent paralysis (Fig. 1K) and swimming rate not different from that of non-paralyzed worms (Fig. 1L). These results indicate that cytosolic ferrous iron is primarily responsible for the observed iron-dependent toxicity.

### 3B. Mitochondrial bioenergetic dysfunction is an early event of iron toxicity

We found that a significant reduction in swimming rate occurred prior to paralysis (Fig 1D & E). Mitochondria supply the bulk of energy needed for cellular function [11, 25, 26]. Therefore, reduced energetic activity could in part be due to an effect of iron on mitochondrial ATP production. To test this hypothesis, we treated worms with and without iron for 3 days, which was the shortest time sufficient to significantly impact mobility (Fig. 1D), but well after impairment of swimming rate (Fig. 1E). Under these conditions, maximum state 3 respiration was significantly reduced (Fig. 2A) in the absence of an effect on state 4 respiration (Fig. 2B), thus resulting in reduced respiratory control ratio (RCR), which is a measure of mitochondrial ATP production (Fig. 2C). Therefore, this iron-dependent reduction in RCR translates to reduced ATP output at a time when worm swimming rate is reduced, and paralysis is increased. We also found that complex I enzyme activity (Fig. 2D) and citrate synthase activity (Fig. 2E) were also significantly reduced in mitochondria isolated from iron-treated worms compared to control worms. Prior studies reported that iron-induced toxicity is mediated in part through increases in ROS production[27, 28]. Therefore, we tested if iron-mediated inhibition of mitochondrial respiratory function resulted in increased mitochondrial ROS generation by monitoring superoxide production with DHE-HPLC[12, 19, 29]. We confirmed increased ROS generation in isolated mitochondria from iron-treated worms compared to that observed for control worms (Fig. 2F). These findings are consistent with iron-induced toxicity resulting in both an inhibition of mitochondrial respiration and an increase in mitochondrial ROS production. High levels of mitochondrial ROS cause oxidative damage of membrane lipid in the form of increased lipid peroxidation (Fig. 1F&G). Having observed impaired mitochondrial bioenergetic function, increased mitochondrial ROS production, and enhanced lipid peroxidation in iron-treated worms, we tested if these effects are due to increased accumulation of mitochondrial ferrous iron. We first measured mitochondrial iron content in control and iron-treated WT and *smf-3* mutant worms. Mitochondrial iron (Fig. 2G) and Ca^2+^ (Fig. 2H) content were significantly higher in iron treated WT worms compared to iron treated *smf-3* worms. Since DMT1 (*smf-3*) also facilitates uptake of other divalent metals[23, 24, 30], we also assessed the impact of *smf-3* mutant on mitochondria Cu^2+^, Mn^2+^ and Zn^2+^ content. While *smf-3* mutant also significantly reduced the mitochondrial content of these divalent metals as expected, mitochondrial levels of these metals were not further impacted by iron exposure (Fig. 2I-K). Overall, the results suggest that ferrous iron toxicity results in mitochondrial dysfunction characterized by reduced aerobic ATP production coupled with increased mitochondrial iron and Ca^2+^ content, ROS production, and lipid peroxidation.

**Figure 2:**
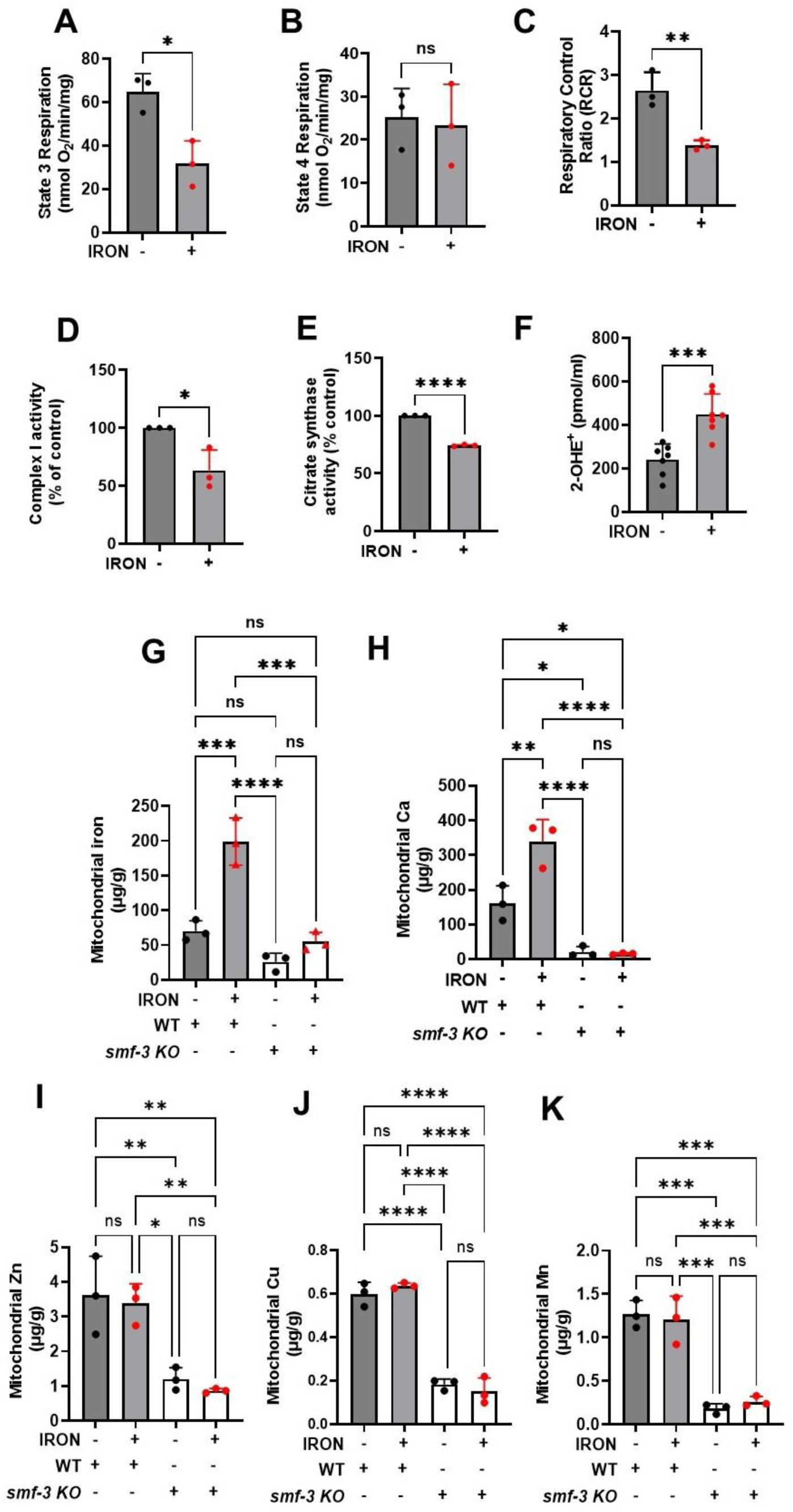
Iron toxicity impairs mitochondrial function and increases ROS production. Synchronized worms (0.25 million/plate) were transferred to plate with or without iron (0 and 35 µM). Mitochondria were isolated after 3 days and quantified for: **A)**. State 3 respiration is the maximum respiratory rates following addition of ATP. Data are mean ±SEM N=3 independent replicates. *p<0.05, Unpaired t test. **B)**. State 4 respiration is the minimum respiratory rates upon depletion of ATP. Data are mean ±SEM N=3 independent replicates. ns not significant, Unpaired t test. **C)**. Respiratory control ratio (RCR) which is the ratio of maximum respiration state 3 over that of minimum respiration state 4. Data are mean ±SEM N=3 independent replicates. **p<0.001, Unpaired t test. **D)**. Iron toxicity reduced mitochondrial complex I enzyme activity. Data are mean ±SEM N=3 independent replicates. *p<0.05, Unpaired t test. **E)**. Iron toxicity reduced citrate synthase activity. Data are mean ±SEM N=3 independent replicates. *p<0.0001, Unpaired t test. **F)**. Iron toxicity increased mitochondrial Superoxide (O_2_^●-^) production. Data are mean ±SEM N=7 independent replicates. ***p=0.0001, Unpaired t test. **G)**. DMT1 (*smf-3*) knock out reduced mitochondrial iron uptake. Mitochondrial iron was measured using ICP-MS in isolated mitochondrial from WT, WT + 35 µM iron, *smf-3* KO, and *smf-3* + 35 µM iron. Data are mean ±SEM, N=3 independent biological replicates. ns not significant, ***p=0.0001, ***p<0.0001, one-way ANOVA, Tukey post hoc test. **H)**. DMT1 (*smf-3*) knock out abolished iron induced increase in mitochondrial calcium uptake. Mitochondrial Ca was measured using ICP-MS in isolated mitochondrial from WT, WT + 35 µM iron, *smf-3* KO, and *smf-3* + 35 µM iron. Data are mean ±SEM, N=3 independent biological replicates. ns not significant, *p<0.05, **p<0.001, ***p<0.0001, one-way ANOVA, Tukey post hoc test. **I)**. Mitochondrial Zn is not impacted by iron toxicity in DMT1 knock out. Mitochondrial Zn was measured using ICP-MS in isolated mitochondrial from WT, WT + 35 µM iron, *smf-3* KO, and *smf-3* + 35 µM iron. Data are mean ±SEM, N=3 independent biological replicates. ns not significant, *p<0.05, **p<0.001, one-way ANOVA, Tukey post hoc test. **J)**. Mitochondrial Cu is not impacted by iron toxicity in DMT1 knock. Mitochondrial Cu was measured using ICP-MS in isolated mitochondrial from WT, WT + 35 µM iron, *smf-3* KO, and *smf-3* + 35 µM iron. Data are mean ±SEM, N=3 independent biological replicates. ns not significant, ***p<0.0001, one-way ANOVA, Tukey post hoc test. **K)**. No effect of iron on mitochondrial Mn in DMT1 knock. Mitochondrial Mn was measured using ICP-MS in isolated mitochondrial from WT, WT + 35 µM iron, *smf-3* KO, and *smf-3* + 35 µM iron. Data are mean ±SEM, N=3 independent biological replicates. ns not significant, ***p=0.0001, one-way ANOVA, Tukey post hoc test.

### 3C. Redox modulation of iron induced oxidative damage

The above results suggest that iron-induced mitochondrial dysfunction results in increased ROS production and oxidative damage. Thus, reducing or enhancing mitochondrial ROS levels should mitigate or exacerbate, respectively, iron mediated toxicity. Indeed, iron-induced paralysis was decreased and swimming rate was increased following treatment with a mitochondrial-targeted antioxidant (Mito-TEMPO) that traps superoxide[12, 31, 32] (Fig. 3A-C). These findings are consistent with iron-induced mitochondrial toxicity resulting from an increase in mitochondrial superoxide levels (Fig. 2F). Superoxide is rapidly dismutated through both non-enzymatic and enzymatic (superoxide dismutase or SOD) means[12]. Manganese (III) Porphyrin (Mn(III)PyP) is an SOD mimetic that converts superoxide to H_2_O_2_[12, 33]. Thus, Mn(III)PyP increases the flux rate of conversion of superoxide to H_2_O_2_, thereby increasing levels of mitochondrial H_2_O_2_. Mn(III)PyP exacerbated iron-induced paralysis and reduced swimming rate (Fig. 3D-F), consistent with toxic effects of increased levels of H_2_O_2_. These results suggest that the species of ROS responsible for the iron mediated toxicity is H_2_O_2_ (and/or an H_2_O_2_ byproduct like hydroxyl radical). Thus, reducing H_2_O_2_ should ameliorate iron-induced toxicity. To test this hypothesis, we used EUK 134, an SOD and catalase mimetic[12, 33, 34], that rapidly converts superoxide to H_2_O_2_ and then water. Treatment with EUK 134 prevented iron-induced worm paralysis and normalized swimming rate (Fig. 3G-I). Similar results were observed by increasing glutathione levels with N-acetyl cysteine (NAC) (Fig. 3J-L). Overall, results in Fig. 3 indicate that the mitochondrial redox environment is a key driver of iron-induced toxicity and that H_2_O_2_ (or its byproducts) is the ROS responsible for iron mediated oxidative damage.

**Figure 3:**
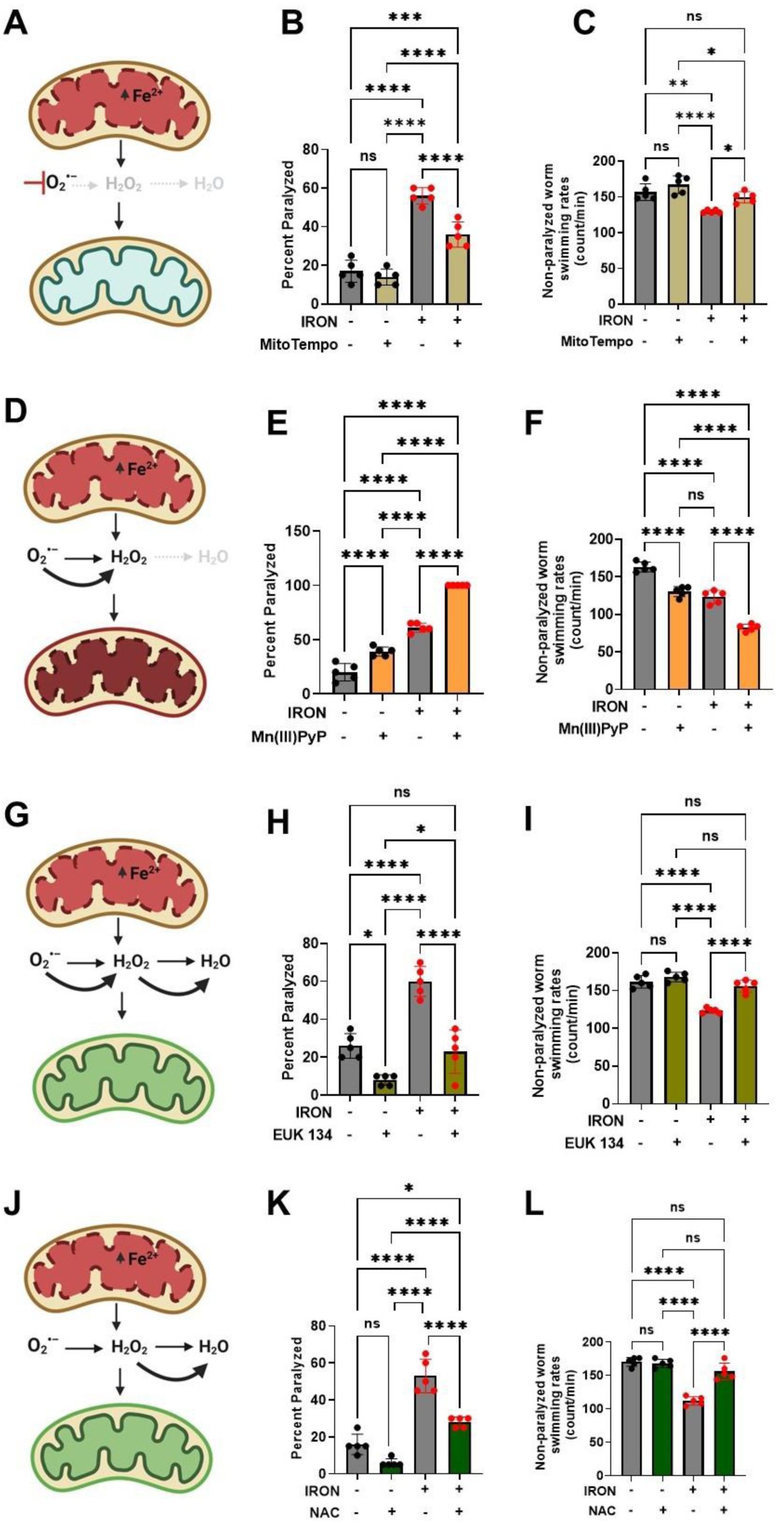
Iron toxicity modulates mitochondrial redox environment. **A)**. Schematic diagram showing specific target of mitoTempo in redox environment. **B)**. MitoTempo ameliorates iron toxicity induced increase in worm paralysis. Staged L4 worms were transferred to plates containing 0 µM iron, 0 µM iron + 10 µM MitoTempo, 35 µM iron and 35 µM iron + 10 µM MitoTempo. Paralysis was scored every 24 h for 5 days. Data are mean ±SEM, N=5 independent biological replicates (where one biological replicate contains 20 worms per plate). ***p=0.0001, ****p<0.0001, one-way ANOVA, Tukey post hoc test. **C)**. MitoTempo restored non-paralyzed worm swimming rates in iron toxicity environment. Staged L4 worms were transferred to plates containing 0 µM iron, 0 µM iron + 10 µM MitoTempo, 35 µM iron and 35 µM iron + 10 µM MitoTempo. Non-paralyzed worms were individually transferred to plate containing 100 µl of buffer. After 30 seconds of equilibration, swimming rates were collected for 15 seconds. Data are mean ±SEM N=5 independent replicates (where 4 independent worm count constitute an N). ns not significant, **p<0.001, ****p<0.0001, one-way ANOVA, Tukey post hoc test. **D)**. Schematic diagram showing specific target of SOD mimetic manganese porphyrin [Mn(III)PyP] in redox environment. **E)**. Mn(III)PyP exacerbates iron toxicity induced increase in worm paralysis. Staged L4 worms were transferred to plates containing 0 µM iron, 0 µM iron + 100 µM Mn(III)PyP, 35 µM iron and 35 µM iron + 100 µM Mn(III)PyP. Paralysis was scored every 24 h for 5 days. Data are mean ±SEM, N=5 independent biological replicates (where one biological replicate contains 20 worms per plate). ****p<0.0001, one-way ANOVA, Tukey post hoc test. **F)**. Mn(III)PyP worsen non-paralyzed worm swimming rates in iron toxicity environment. Staged L4 worms were transferred to plates containing 0 µM iron, 0 µM iron + 100 µM Mn(III)PyP, 35 µM iron and 35 µM iron + 100 µM Mn(III)PyP. Non-paralyzed worms were individually transferred to plate containing 100 µl of buffer. After 30 seconds of equilibration, swimming rates were collected for 15 seconds. Data are mean ±SEM N=5 independent replicates (where 4 independent worm count constitute an N). ****p<0.0001, one-way ANOVA, Tukey post hoc test. **G)**. Schematic diagram showing specific target of EUK 134 (SOD and Catalase mimetic) in mitochondrial redox environment. **H)**. EUK 134 protects against iron-induced toxicity induced in worm paralysis. Staged L4 worms were transferred to plates containing 0 µM iron, 0 µM iron + 100 µM EUK 134, 35 µM iron and 35 µM iron + 100 µM EUK 134. Paralysis was scored every 24 h for 5 days. Data are mean ±SEM, N=5 independent biological replicates (where one biological replicate contains 20 worms per plate). ns not significant ***p=0.0001, ****p<0.0001, one-way ANOVA, Tukey post hoc test. **I)**. EUK 134 restored non-paralyzed worm swimming rates in iron toxicity environment. Staged L4 worms were transferred to plates containing 0 µM iron, 0 µM iron + 100 µM EUK 134, 35 µM iron and 35 µM iron + 100 µM EUK 134. Non-paralyzed worms were individually transferred to plate containing 100 µl of buffer. After 30 seconds of equilibration, swimming rates were collected for 15 seconds. Data are mean ±SEM N=5 independent replicates (where 4 independent worm count constitute an N). ns not significant, **p<0.001, ****p<0.0001, one-way ANOVA, Tukey post hoc test. **J)**. Schematic diagram showing specific target of N-Acetyl Cysteine (NAC) in mitochondrial redox environment. **K)**. NAC protects against iron toxicity induced increase in worm paralysis. Staged L4 worms were transferred to plates containing 0 µM iron, 0 µM iron + 2.5 mM NAC, 35 µM iron and 35 µM iron + 2.5 mM NAC. Paralysis was scored every 24 h for 5 days. Data are mean ±SEM, N=5 independent biological replicates (where one biological replicate contains 20 worms per plate). *p<0.05, ***p=0.0001, ****p<0.0001, one-way ANOVA, Tukey post hoc test. **L)**. NAC restored non-paralyzed worm swimming rates in iron toxicity environment. Staged L4 worms were transferred to plates containing 0 µM iron, 0 µM iron + 2.5 mM NAC, 35 µM iron and 35 µM iron + 2.5 mM NAC. Non-paralyzed worms were individually transferred to plate containing 100 µl of buffer. After 30 seconds of equilibration, swimming rates were collected for 15 seconds. Data are mean ±SEM N=5 independent replicates (where 4 independent worm count constitute an N). ns not significant, ****p<0.0001, one-way ANOVA, Tukey post hoc test

### 3D. Ferroptosis mediates iron induced oxidative damage

Our findings that iron-induced toxicity causes increased levels of ROS (Fig. 2F) and lipid peroxidation (Fig. 1F&G) are consistent with a potential role for ferroptosis, or iron-induced cell death through ROS mediated lipid peroxidation[3, 13]. Therefore, we used canonical ferroptosis modulators (Ferrostatin-1 and RSL3) to assess the involvement of ferroptosis in the observed iron-induced toxicity. Ferrostatin-1 (Fer-1), a small molecule that inhibits lipid peroxidation by scavenging alkoxyl radicals (Fig. 4A), which subsequently leads to reduction in ferroptosis[3, 35–37], abolished both the iron mediated increase in paralysis (Fig. 4B) and decrease in swimming rate (Fig. 4C), consistent with a role for ferroptosis. To more critically assess the role of ferroptosis, we used RSL3, a potent inhibitor of GPX4[3, 38], a selenoenzyme with phospholipid hydroperoxidase activity that converts PUFA-OOH to PUFA-OH (Fig. 4A), thus neutralizing phospholipid oxidative damage[3, 39, 40]. Unlike the mammalian GPX4, *C. elegans gpx1* does not contain selenium, but possesses phospholipid hydroperoxidase activity like the mammalian GPX4[41, 42]. In addition, not all mammalian selenium containing glutathione peroxidases possess phospholipid hydroperoxidase activity[43]. Thus, GPX4 PUFA oxidation activity could be mediated through phospholipid hydroperoxidase. We tested if the *C. elegans* glutathione peroxidase (*gpx1)* exhibits phospholipid hydroperoxidase activity in response to RSL3 as is observed for mammalian GPX4 [3, 39, 40]. RSL3 alone induced paralysis (Fig. 4D) and reduced swimming rates of non-paralyzed worms (Fig. 4E). Further, this RSL3 exacerbation of iron-induced toxicity was abolished by Fer-1 (Fig. 4D & E). Taken together, these data suggest the involvement of ferroptosis in iron mediated pathology and support the idea that iron-induced toxicity is mitigated by interventions that inhibit ferroptosis.

**Figure 4:**
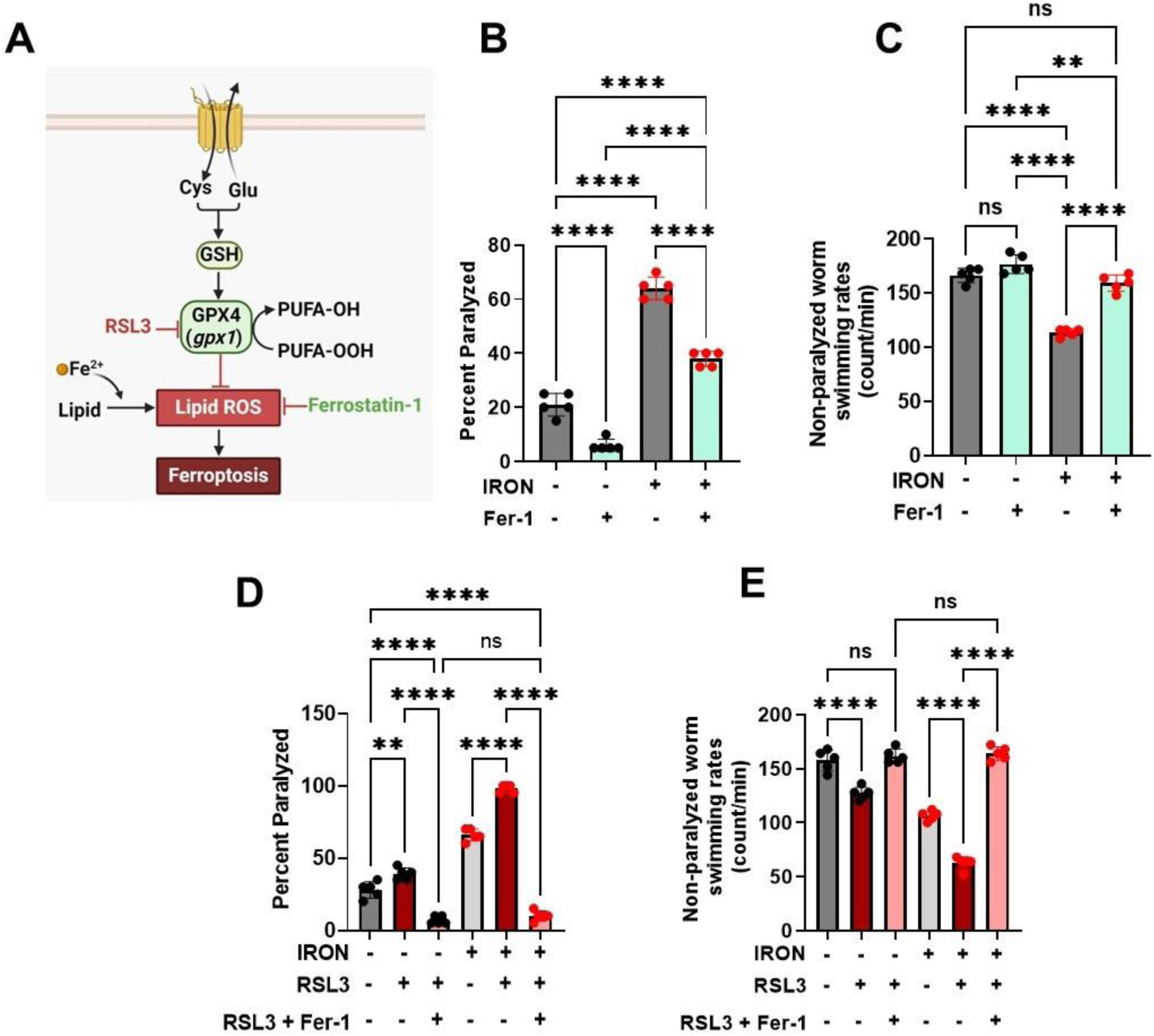
Ferroptosis modulates iron induced toxicity. **A)**. Schematic diagram showing side of ferroptotic drug target regulating iron toxicity. **B)**. Ferrostatin-1 (Fer-1) ameliorates iron-induced worm paralysis. Staged L4 worms were transferred to plates containing 0 µM iron, 0 µM iron + 5 µM Fer-1, 35 µM iron and 35 µM iron + 5 µM Fer-1. Paralysis was scored every 24 h for 5 days. Data are mean ±SEM, N=5 independent biological replicates (where one biological replicate contains 20 worms per plate). ****p<0.0001, one-way ANOVA, Tukey post hoc test. **C)**. Ferrostatin-1 restored non-paralyzed worm swimming rates in iron toxicity environment. Staged L4 worms were transferred to plates containing 0 µM iron, 0 µM iron + 5 µM Fer-1, 35 µM iron and 35 µM iron + 5 µM Fer-1. Non-paralyzed worms were individually transferred to plate containing 100 µl of buffer. After 30 seconds of equilibration, swimming rates were collected for 15 seconds. Data are mean ±SEM N=5 independent replicates (where 4 independent worm count constitute an N). ns not significant, ****p<0.0001, one-way ANOVA, Tukey post hoc test. **D)**. RSL3 exacerbates iron toxicity induced increase in worm paralysis. Staged L4 worms were transferred to plates containing 0 µM iron, 0 µM iron + 5 nM RSL3, 0 µM iron + 5 nM RSL3 + 5 µM Fer-1, 35 µM iron, 35 µM iron + 5 nM RSL3, and 35 µM iron + 5 nM RSL3 + 5 µM Fer-1. Paralysis was scored every 24 h for 5 days. Data are mean ±SEM, N=5 independent biological replicates (where one biological replicate contains 20 worms per plate). ****p<0.0001, one-way ANOVA, Tukey post hoc test. **E)**. RSL3 worsen non-paralyzed worm swimming rates in iron toxicity environment. Staged L4 worms were transferred to plates containing 0 µM iron, 0 µM iron + 5 nM RSL3, 0 µM iron + 5 nM RSL3 + 5 µM Fer-1, 35 µM iron, 35 µM iron + 5 nM RSL3, and 35 µM iron + 5 nM RSL3 + 5 µM Fer-1. Non-paralyzed worms were individually transferred to plate containing 100 µl of buffer. After 30 seconds of equilibration, swimming rates were collected for 15 seconds. Data are mean ±SEM N=5 independent replicates (where 4 independent worm count constitute an N). ****p<0.0001, one-way ANOVA, Tukey post hoc test

### 3E. Worms expressing neuronal Aβ exhibit an enhanced ROS- and ferroptosis- dependent phenotype

In *C. elegans*, movement disorder (i.e., paralysis) is a readout for decline in neuronal function, which is similarly associated with most neurodegenerative disorders including AD[17, 44]. To investigate the impact of iron toxicity on worm models of AD, we exposed WT and pan-neuronal Aβ worms (worms expressing human amyloid beta in all neurons)[44] to iron. Neuronal Aβ worms exhibited greater paralysis and slower swimming rates compared to WT worms under both control conditions and following exposure to 35 µM iron (Fig. 5A & B). The enhanced control, as well as iron induced, paralysis and reduced swimming rate of neuronal Aβ worms were all blocked by treatment with MitoTempo, EUK 134, and NAC, but further enhanced by Mn(III)PyP (Fig. 5C & D). Next, we probed the impact of ferroptosis modulators on the enhanced paralysis and reduced swimming rate phenotype of neuronal Aβ worms. Similar to that observed for modulators of ROS, activator of ferroptosis (RSL3) potentiated, while a ferroptosis inhibitor (Fer-1) reduced, the enhanced paralysis/swimming rate phenotype of neuronal Aβ worms (Fig. 5E & F). These results indicate that ferroptosis plays an important role in the increased paralysis and reduced swimming rate of neuronal Aβ worms, thus supporting the notion that ferroptosis represents a potential target for AD intervention.

**Figure 5:**
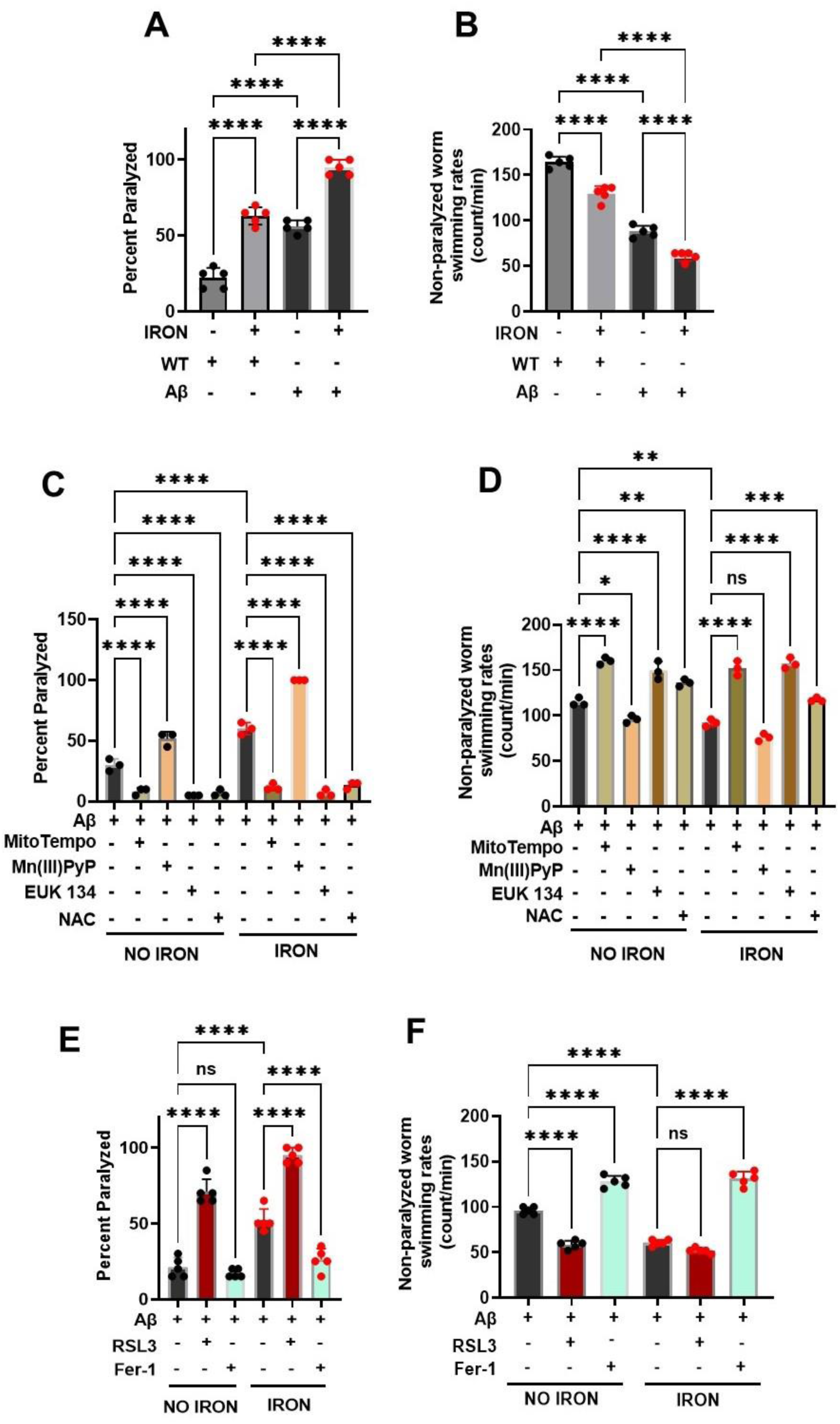
Iron toxicity exacerbates ROS and ferroptosis dependent phenotypes in neuronal Aβ pathology: **A)** Iron toxicity potentiate Aβ increased worm paralyses. Staged L4 worms (WT and neuronal Aβ) were transferred to plates containing iron (0 or 35 µM). Paralysis was scored every 24 h for 5 days. Data are mean ±SEM, N=5 independent biological replicates (where one biological replicate contains 20 worms per plate). ns not significant, ****p<0.0001, one-way ANOVA, Tukey post hoc test. **B)** Iron toxicity worsen Aβ decreased in worm swimming rates. Staged L4 worms (WT and neuronal Aβ) were transferred to plate containing iron (0 or 35 µM). Worms were then transferred every 24 h for 5 days. Non-paralyzed worms were individually transferred to plate containing 100 µl of buffer. After 30 seconds of equilibration, swimming rates were collected for 15 seconds. Data are mean ±SEM N=5 independent replicates (where 4 independent worm count constitute an N). ****p<0.0001, one-way ANOVA, Tukey post hoc test. **C)** Iron toxicity induced ROS potentiates Aβ paralyses. Staged L4 worms neuronal Aβ were transferred to plates containing iron (0 or 35 µM) with oxidative stress modulators 10 µM MitoTempo, 100 µM Mn(III)PyP, 100 µM EUK 134 and 2.5 mM NAC. Paralysis was scored every 24 h for 3 days. Data are mean ±SEM, N=3 independent biological replicates (where one biological replicate contains 20 worms per plate). ****p<0.0001, one-way ANOVA, Tukey post hoc test. **D)** Impact of iron-induced ROS on neuronal Aβ worm swimming rates. Staged L4 worms neuronal Aβ were transferred to plates containing iron (0 or 35 µM) with oxidative stress modulators 10 µM MitoTempo, 100 µM Mn(III)PyP, 100 µM EUK 134 and 2.5 mM NAC. Worms were then transferred every 24 h for 3 days. Non-paralyzed worms were individually transferred to plate containing 100 µl of buffer. After 30 seconds of equilibration, swimming rates were collected for 15 seconds. Data are mean ±SEM N=3 independent replicates (where 4 independent worm count constitute an N). ns not significant, *p<0.05, **p<0.001, ***p=0.0001, ****p<0.0001, one-way ANOVA, Tukey post hoc test. **E)** Ferroptosis regulates iron-induced paralysis in neuronal Aβ pathology. Staged L4 worms neuronal Aβ were transferred to plates containing iron (0 or 35 µM) ferroptosis modulators 5 nM RSL3 and 5 µM Fer-1. Paralysis was scored every 24 h for 3 days. Data are mean ±SEM, N=5 independent biological replicates (where one biological replicate contains 20 worms per plate). ns not significant, ****p<0.0001, one-way ANOVA, Tukey post hoc test. **F)** Ferroptosis drives iron-induced slow swimming of neuronal Aβ worms. Staged L4 worms neuronal Aβ were transferred to plates containing iron (0 or 35 µM) ferroptosis modulators 5 nM RSL3 and 5 µM Fer-1. Worms were then transferred every 24 h for 3 days. Non-paralyzed worms were individually transferred to plate containing 100 µl of buffer. After 30 seconds of equilibration, swimming rates were collected for 15 seconds. Data are mean ±SEM N=5 independent replicates (where 4 independent worm count constitute an N). ns not significant, ****p<0.0001, one-way ANOVA, Tukey post hoc test.

### 3F. Neuronal Aβ worms exhibit enhanced DMT1-dependent iron sensitivity

To test if neuronal Aβ worms are more sensitive to iron toxicity, we exposed worms to 8.75 µM iron, a concentration of iron that is not toxic to WT worms. We found that while exposure for 5 days to 8.75 µM iron had no effect on WT worms, neuronal Aβ worm paralysis was increased, and the swimming rate of non-paralyzed worms was reduced (Fig. 6A & B). Importantly, total worm iron accumulation under these conditions was not different between WT and neuronal Aβ worms (Fig. 6C), suggesting that both WT and neuronal Aβ worms exhibited a similar tissue iron burden. Together, these data indicate that neuronal Aβ worms are more sensitive to iron toxicity relative to WT worms. We then tested if knockout of DMT1 (*smf-3*), which prevented iron-induced toxicity in WT worms, prevented the enhanced basal and iron-induced phenotype of neuronal Aβ worms. DMT1 (*smf-3*) mutant significantly reduced paralysis and increased swimming rate of neuronal Aβ worms under control conditions and also reduced the degree of iron-induced paralysis and swimming rate of these worms (Fig. 6D & E). Importantly, *smf-3* knockout not only significantly reduced the effects of Aβ overexpression on work activity (i.e., swimming rate), it restored these measures in Aβ worms to WT levels. Overall, results in Fig. 6 demonstrate that genetic disruption of DMT1 (*smf-3*) function mitigates both basal and iron-induced pathologies of neuronal Aβ worms, and thus, represent a new potential therapeutic target for AD.

**Figure 6:**
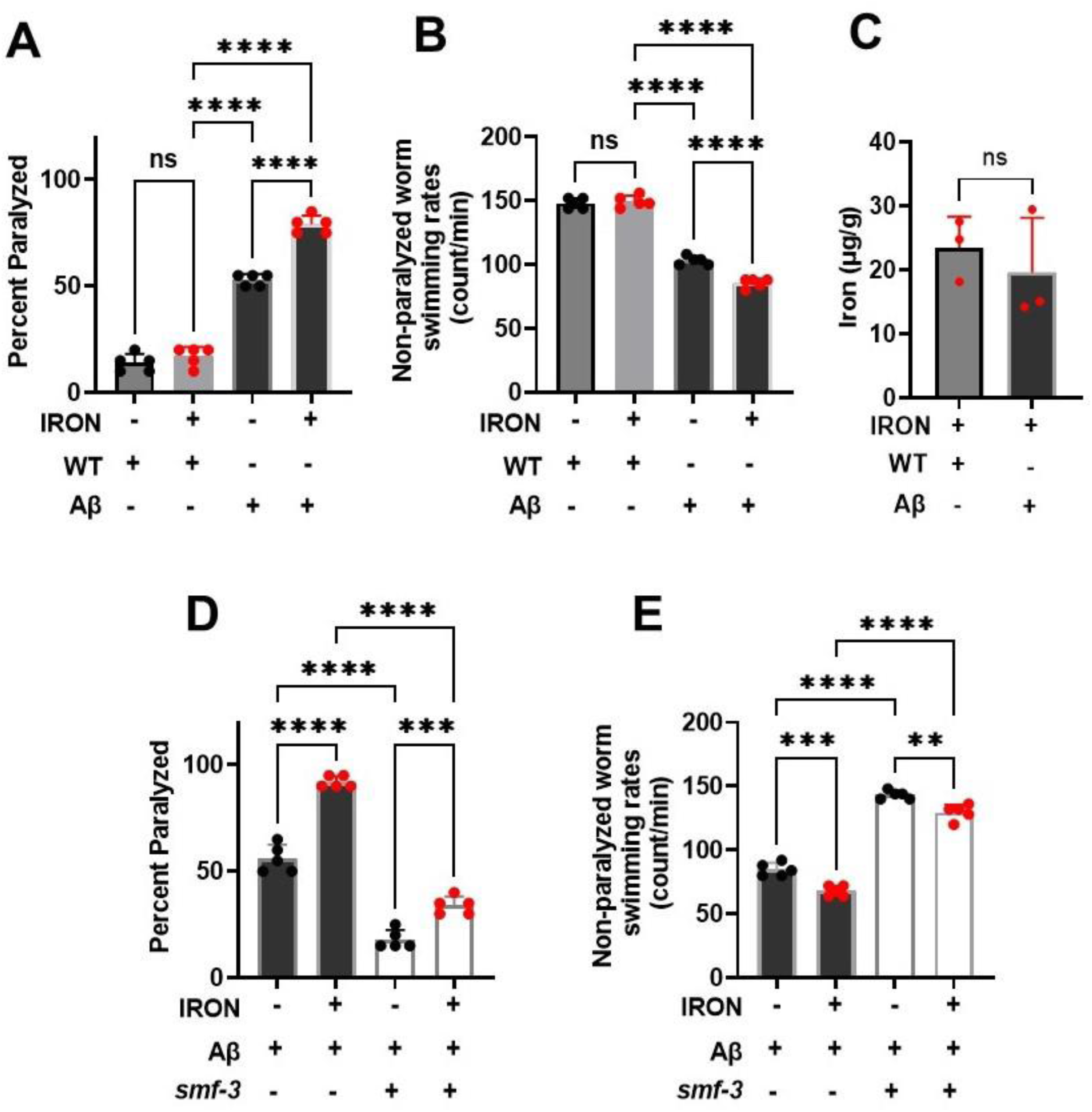
DMT1(*smf-3*) KO protects against neuronal Aβ pathology: Neuronal Aβ worms exhibit increase sensitivity to iron toxicity than WT. **A)** Paralysis: Staged L4 worms (WT and neuronal Aβ) were transferred to plates containing iron (0 or 8.75 µM). Paralysis was scored every 24 h for 5 days. Data are mean ±SEM, N=5 independent biological replicates (where one biological replicate contains 20 worms per plate). ns not significant, ****p<0.0001, one-way ANOVA, Tukey post hoc test. **B)** Non-paralyzed worm swimming rates. Staged L4 worms (WT and neuronal Aβ) were transferred to plate containing iron (0 or 8.75 µM). Worms were then transferred every 24 h for 5 days. Non-paralyzed worms were individually transferred to plate containing 100 µl of buffer. After 30 seconds of equilibration, swimming rates were collected for 15 seconds. Data are mean ±SEM N=5 independent replicates (where 4 independent worm count constitute an N). ns not significant, ****p<0.0001, one-way ANOVA, Tukey post hoc test. **C)** Neuronal Aβ iron sensitivity not mediated by iron burden. Tissue iron burden was measured in WT and Aβ treated with 8.75 µM iron for 5 days using ICP-MS. Data are mean ±SEM, N=3 independent biological replicates. ns not significant, Unpaired t test. **D)**. Knock out of *smf-3* in neuronal Aβ worms abolished paralysis. Staged L4 worms (neuronal Aβ and neuronal Aβ+ *smf-3* KO) were transferred to plates containing iron (0 or 35 µM). Paralysis was scored every 24 h for 5 days. Data are mean ±SEM, N=5 independent biological replicates (where one biological replicate contains 20 worms per plate). ***p=0.0001, ****p<0.0001, one-way ANOVA, Tukey post hoc test. **E)** *smf-3* KO in neuronal Aβ worms protects against Aβ decreased swimming rate. Staged L4 worms (neuronal Aβ and neuronal Aβ+ *smf-3* KO) were transferred to plates containing iron (0 or 35 µM). Worms were then transferred every 24 h for 5 days. Non-paralyzed worms were individually transferred to plate containing 100 µl of buffer. After 30 seconds of equilibration, swimming rates were collected for 15 seconds. Data are mean ±SEM N=5 independent replicates (where 4 independent worm count constitute an N). **p<0.001, ***p=0.0001, ****p<0.0001, one-way ANOVA, Tukey post hoc test.

## 4. Discussion

As organisms age, the capacity of iron regulatory proteins to properly sequester and control cellular iron weakens resulting in age-related pathologies that contribute to neurodegeneration such as that observed in AD[3, 7, 8]. Here we document a similar observation in *C. elegans*, wherein iron pathology is further exacerbated by neuronal Aβ. Thus, regulation of cellular iron levels is a critical step in iron-dependent pathologies. Our study focused on two related phenotypes (swimming rate and paralysis) with distinct iron pathology. For example, paralysis (an inability to move) correlates with a decline in neuronal function. We demonstrated that enhanced iron accumulation occurs prior to progressive paralysis, which mirrors the loss of neuronal function in AD that worsens with age[45, 46]. In contrast to paralysis, swimming rate reflects energetic output and reduced swimming rate occurs earlier than loss of neuronal function (paralysis) in WT worms. Reduced swimming rate could also be due to a partial reduction in neuronal function while paralysis reflects progression to complete loss of neuronal function. A decrease in mitochondrial energy production is a likely explanation for the observed iron-induced reduction in swimming rate. Therefore, therapies designed to target mitochondrial iron could alleviate this decline in mitochondrial function. Yet the underlying molecular mechanisms of mitochondrial iron toxicity in AD remain elusive. We propose that ferrous iron inhibition of mitochondrial respiration results in increased mitochondrial ROS generation that ultimately leads to modifications of ETC proteins and increased oxidative damage (e.g., lipid peroxidation).

We found that ferrous iron-induced mitochondrial dysfunction drives an increase in oxidative damage in both WT and neuronal Aβ worms. Others similarly reported that iron exposure promotes oxidative damage[47, 48]. However, precisely how ferrous iron modulates the mitochondrial redox environment to enhance oxidative damage remains unclear. We demonstrate that ferrous iron exposure enhanced both mitochondrial ROS production and lipid peroxidation, consistent with changes in the mitochondrial redox environment playing a significant role in ferrous iron-induced pathology[49]. In support of this assertion, interventions designed to either exacerbate or abolish mitochondrial ROS levels increased or decreased, respectively, the ferrous iron induced pathology of both WT and neuronal Aβ worms. Similar to redox changes, we also found that ferrous iron-induced toxicity in WT and neuronal Aβ worms was increased or decreased by ferroptosis inducers and inhibitors, respectively. Overall, our studies demonstrate that ferrous iron-induced toxicity in WT and neuronal Aβ worms is mediated by mitochondrial dysfunction leading to reduced aerobic ATP production that occurs in concert with increased oxidative stress, lipid peroxidation, and ferroptosis.

Brains of AD patients exhibit high levels of iron relative to that of age-matched non-AD individuals[50, 51]. However, it is unclear if increased brain iron levels drive AD pathology. In addition, it is unknown whether AD patients are more sensitive to iron-induced oxidative stress than non-AD individuals. We found that neuronal Aβ worms exhibited increased pathology under control conditions, greater pathology following exposure to a moderate level of ferrous iron (Fig. 5) and increased sensitivity to low levels of ferrous iron that do not impact age-matched WT worms (Fig. 6). Thus, our studies demonstrate that neuronal Aβ worms exhibit both a greater responsiveness and higher sensitivity to ferrous iron. We further found that the increase in ferrous iron-induced sensitivity of neuronal Aβ worms is not due to increased ferrous iron uptake (Fig. 6C), but rather due to an increase in susceptibility to ferrous iron-induced stress.

DMT1 (*smf-3*) facilitates cellular and mitochondrial uptake of ferrous iron[30, 52, 53]. Thus, we tested if DMT1 (*smf-3*) knockout in WT and neuronal Aβ worms would provide protection against ferrous iron-induced toxicity. Importantly, DMT1 (*smf-3*) knockout not only mitigated toxicity induced by ferrous iron exposure in both WT and neuronal Aβ worms, but also reduced the basal AD pathology of neuronal Aβ worms. As organisms age, their ability to properly handle ferrous iron weakens[3, 7, 8], which explains the beneficial effects DMT1 (*smf-3*) knockout of neuronal Aβ worms in the absence of iron exposure. Precisely how DMT1 (*smf-3*) deficiency mediates this protection (e.g. via reduction of cellular iron uptake and/or reduction in mitochondrial iron accumulation) will require further study. Our study revealed that increased sensitivity of neuronal Aβ worms to properly handle ferrous iron results in pathology and that disrupting ferrous iron transport in this model of AD provides protection against iron-induced oxidative stress and ferroptotic cell death. Overall, our findings suggest that the energetic imbalance resulting from DMT1 (*smf-3*)-dependent ferroptosis may represent an early event of AD pathogenesis.

## Authors Contributions

JOO conceived and designed the study, JOO, WP, KBC carried out the experiments, JOO, APW, RTD, BPL, MKO supervised the study, JOO wrote the manuscript. All authors reviewed, edited, and approved the final manuscript.

## Acknowledgements

We thank the Center for Advanced Light Microscopy and Nanoscopy (CALMN) of the University of Rochester for their technical assistance. We are grateful to Thomas Scrimale of the Environmental Health Sciences Core (EHSC) for the Elemental analysis. Most of the strains used in this study were obtained from the *Caenorhabditis* Genetics Center (CGC), which is supported by NIH. All schematic illustrations and graphical image were generated with BioRender. We appreciate the rigorous discussions and input from the Mitochondrial Research & Innovation Group at University of Rochester Medical Center, the Western New York Worm Group, and the Journal Club of Pharmacology and Physiology Department.

## Data availability

All data are available upon request

